# No limit yet to phenological plasticity of reproductive timing in a high-elevation hibernator experiencing earlier springs

**DOI:** 10.1101/2025.11.19.688555

**Authors:** Caitlin P. Wells, Dirk H. Van Vuren

## Abstract

Plasticity in the timing of reproduction may enable species to adapt to changing climates and match reproductive demands to shifting peaks of energy availability. At high elevations, hibernating mammals are often facing drier winters and earlier snowmelt, shifting the onset of the vegetative growing season. We expect substantial plasticity in reproductive phenology at these sites because inter-annual variation in snowmelt date is typically high. However, whether species are reaching limits of phenological plasticity with ongoing directional change has rarely been assessed for vertebrate populations; identification of phenological limits is important to predict further adaptive capacity. Here, we use 30 years of litter weaning dates of golden-mantled ground squirrels (*Callospermophilus lateralis*) from the Rocky Mountains in Colorado, USA, to assess individual plasticity in reproductive phenology, population trends over time, and whether the population has reached a limit of plasticity. On average, litter weaning dates advanced by 2.3days/decade over the study period. We found that individual reproductive timing was quite plastic and exhibited near-zero heritability. At the population level, a linear relationship between average date of litter weaning and date of snowmelt suggests further potential for phenological adaptation to earlier springs. However, litters emerge later in relation to vegetative phenology in early springs, generating a potential mismatch between peak vegetative abundance and energetic demands of mothers and pups.

## Introduction

Changes in phenology – the timing of annual events such as migration and reproduction – is one of the clearest responses of animals to ongoing climatic change (Visser & Both 2005, Parmesan 2006, Cohen et al. 2018, Post et al. 2018, Inouye 2022). Changing phenology to match environmental change is presumably adaptive, as maintaining synchrony with interacting species should increase survival and reproduction (Renner & Zohner 2018). However, observations of phenological change are often at the species or population level, making it difficult to link patterns to mechanisms at the individual level (Chmura et al. 2019). Importantly, we know for few species whether phenological changes are a product of microevolutionary change (e.g. Balanya et al. 2006), phenotypic plasticity (e.g. Charmantier 2008), or both (e.g. Crozier and Hutchings 2014, Bonnet et al. 2019, Moiron et al. 2024), and differentiating between the two requires individual-level data (Charmantier & Gienapp 2014). Identifying the plastic or genetic bases for changes in phenology are essential to predict short-versus long-term responses and the persistence of species experiencing continued changes in temperature, precipitation, and seasonality (Merilä & Hendry 2014).

Hibernation is a phenological event that might be particularly sensitive to changes in climate (Wells et al. 2022, Findlay-Robinson et al. 2023). Hibernating species use seasonal lowering of metabolic rates (i.e. torpor) to survive periods of food scarcity – typically during the cold winter for temperate species – timing emergence from hibernation to coincide with new food availability (Geiser and Ruf 1995). As the lowered metabolic rates of hibernation impede gestation, small-bodied hibernators typically time reproduction during the active euthermic period via delayed implantation of embryos, arrested gestation, or ovulation within days of emergence in spring (Oxberry 1979, Vasilieva et al. 2024). In ground squirrels, reproduction is initiated immediately after emergence from hibernation in spring in response to temporal constraints of a short vegetative growing season: to survive, adult females producing a litter must also have time after weaning to regain sufficient fat stores to fuel their hibernation (Armitage 2014). Emergence phenology has been relatively well-studied for hibernating species and reliably shown to advance with warmer temperatures and shorter winters (Wells et al. 2022). However, reproductive phenology is substantially less studied, and may become decoupled from emergence when individual condition is poor (Millesi et al. 1999).

If reproductive timing (phenology) is genetically based, it should show repeatability and heritability. The magnitude of heritability will allow us to estimate if timing can evolve in response to changing ecological conditions, and how quickly, as a trait with a larger heritability should respond more strongly to selection (Wilson 2010). Alternatively, if there is little or no repeatability or heritability in phenology, animals may still adapt via individual plasticity in reproductive timing. For example, if environmental cues of optimal breeding conditions are reliable, animals may be able use those to initiate reproduction (Wingfield et al. 1992). A third possibility is intermediate between these two extremes, wherein if reproductive timing is set during development, it may show individual repeatability but no heritability; this would allow individuals to adapt to ecological conditions during their first reproductive bout, but would not extend to subsequent generations and could be a constraint if conditions differ in subsequent reproductive bouts over the course of the individual’s lifetime. Additionally, reproductive timing may vary across the lifespan due to internal cues, presumably based on changes in individual condition: mid-aged individuals often show earlier phenology than first-time breeders (Bonamour et al. 2020, Oosthuizen et al. 2023), but sometimes not (St. Lawrence et al. 2024).

Reproductive phenology may be controlled by a combination of endogenous and environmental cues. Endogenous circannual cycles have been documented in a range of taxa, from plants to arthropods, fish, reptiles, birds, and mammals (Gwinner 2012). These annual rhythms are termed “circannual” because in the absence of environmental cues, the endogenous rhythm has a period less than 12 months. The hibernation cycle of golden-mantled ground squirrels (*Callospermophilus lateralis*) was the first experimental demonstration of circannual periodicity, with captive individuals maintaining an ∼11 month cycle under constant photoperiod and temperature conditions (Pengelley 1957, 1963). Further work identified the interrelatedness of the hibernation, metabolic, and reproductive cycles in this species, with the maintenance of clear phase relationships between them (e.g. 3 months from peak body mass to peak testicular development for males, Kenagy 1980). However, integration of a hierarchical set of cues is expected to “fine tune” the circannual cycle: while the broad rhythms are controlled endogenously, proximate cues of upcoming food availability (e.g. temperature or precipitation) are expected to entrain the circannual cycle to the 12-month cycle of environmental conditions (Gwinner 2012). Adjustment of endogenous circannual cycles of reproduction to environmental conditions is essential to adapt to changing climates and allow hibernating species to persist (Williams et al. 2014; Aubry & Williams 2022)

Another important question about phenological adaptation to future conditions is whether animals have already reached a threshold of plasticity. A linear response to a changing component of climate, such as temperature or spring snowmelt, suggests that additional plasticity at ranges outside of current conditions is possible (Iler et al. 2013). A nonlinear response, however, showing a flattening of phenological response at high temperatures or early springs, suggests that a threshold of plasticity has been reached. At high elevations, advancement of flowering time in response to spring conditions appears to have reached a threshold for some plant species but not others (Iler et al. 2013). However, climatic thresholds of phenological plasticity have not been evaluated for many vertebrate species (Vedder et al. 2013).

Finally, the individual fitness consequences of plasticity in reproductive timing likely depend on the phenology of other taxa. Differences in the rate at which interacting species change their phenology in response to environmental conditions is known as phenological mismatch or trophic asynchrony (Stenseth & Mysterud 2002, Renner & Zohner 2018). Plasticity in reproductive phenology may allow adult females to track environmental changes in seasonal resource availability, i.e. to “match” changes in phenology of other trophic levels. Alternatively, less plasticity than that of key resources – i.e. “mis-match” – could reduce body mass gain for reproducing females and their offspring. In terrestrial systems lower trophic levels are generally more plastic than higher trophic levels, meaning consumers are often shifting less than the producers they rely on (Thackeray et al. 2010). Despite several classic studies on consequences of mismatches in phenology in birds (Visser et al. 1998, Franks et al. 2017), relatively little work has been published on mammals (e.g. Plard et al. 2014, Renaud et al. 2022; Kharouba & Wolkovich 2023). As temperatures are warming more rapidly at high-elevations (Pepin et al. 2022), evaluating phenological mismatch for hibernating mammals there would point to whether plasticity of primary consumers can be sufficient to match changes in highly seasonal environments, and indicate if the phenomenon is broadly applicable (Kharouba & Wolkovich 2023).

Golden-mantled ground squirrels (hereafter, “GMGS”) are small bodied (200-300g), diurnal mammals, which hibernate over the Northern hemisphere winter (McKeever 1964). They are predominantly herbivorous, and at high elevations rely on forbs and grasses that begin growth once snowpack melts in spring (Carleton 1966). In GMGS, the seasonal timing of reproduction is extremely important for survival and development of pups, as females weaned relatively later in the summer show reduced growth rates and delayed reproductive maturity (from age 1 to age 2), presumably due to reduced vegetation quality or quantity (Wells & Van Vuren 2018, Howland et al. 2024). Additionally, a focus on reproductive timing of females is important because ground squirrels typically exhibit sex-specific phenology (e.g. Michener 1983, Healy et al. 2012): adult males typically emerge from hibernation well before females, and phenology estimates based on first-of-year sightings (e.g. Inouye et al. 2000) are therefore likely male-specific. Moreover, phenological plasticity may differ between the sexes, with females of some species showing more sensitivity than males to changes in spring cues (Williams et al. 2017, Chmura et al. 2023), generating the potential for sexual phenological mismatch (Nakazawa et al. 2023).

In this study, we ask first if reproductive phenology (i.e. date of litter weaning) of female golden-mantled ground squirrels is tracking environmental change. Second, we evaluate if the relationship between date of litter weaning and environmental change is linear or non-linear, for evidence that the population is reaching a limit on plasticity in reproductive phenology. Third, we ask if there is any individual repeatability in date of litter weaning in response to endogenous or environmental cues. Fourth, we calculate the heritability of date of litter weaning for evidence of genetic versus plastic contributions to phenological shifts. Last, we evaluate the relationship between date of litter weaning and vegetation phenology for evidence of a trophic match or mismatch in years with earlier springs.

## Methods

### Study system

We studied one population of golden-mantled ground squirrels at the Rocky Mountain Biological Laboratory (RMBL) in the East River Valley of Colorado, USA (38^°^ 58^’^ N, 106^°^ 59^’^ W, 2900m) from 1990-2023. GMGS at this elevation are active above ground from c. May – September (Wells & Van Vuren 2017). The 13-hectare study site is predominantly open subalpine meadow, interspersed with stands of willow (*Salix* sp.), spruce (*Picea* sp.), and aspen (*Populus tremuloides*); GMGS here tend to avoid dense vegetation and occupy areas of dry meadow and rocky slopes (Aliperti et al. 2022).

We tracked the demography of the population by uniquely marking individuals with numbered Monel fingering eartags (National Band & Tag Company, Newport, KY) for permanent identification, and black dye-marks (Nyanzol-D dye) on the dorsal pelage for re-sighting from a distance. We censused the population each year in early June, recapturing uniquely marked individuals from the previous year, and marking new immigrants. We captured squirrels with live traps (Tomahawk, Model 201) baited with sunflower seeds and peanut butter. We aimed to capture individuals once per month during the summer to track reproductive status and body condition (Wells et al. 2019). At each capture we recorded anogenital distance (mm), body mass (g), and reproductive status (nipple development and features). Most adult females 2 years or older are reproductive each year, while typically half of yearling females become reproductive (Wells & Van Vuren 2018). GMGS pups are born underground after 28 days gestation (Cameron 1967), and remain in their natal burrow until weaning at 30 days of age (Phillips 1981).

We identified female home ranges by regular re-sighting of individuals within the study area. GMGS are classified as asocial (Michener 1983, Armitage 1981), and reproductive adult females exhibit only moderate spatial overlap with other adult females (Jesmer et al. 2011, Aliperti 2020). In daily surveys, we identified individual locations using an alphanumeric grid system of 7mX7m squares (Aliperti 2020), and noted areas of core activity (Wells & Van Vuren 2017). We also noted any litter-directed behaviors such as gathering and carrying dry grass to a burrow as nesting material, or moving pre-weaned pups from one burrow to another (Aliperti et al. 2021); when possible to do without disturbing the female, we followed her to the natal burrow and noted its location to check frequently on subsequent days.

Once an adult female was identified as reproductive (i.e. exhibited swollen nipples), and lactating (i.e. fur around nipples was wet or chewed), we searched her home range daily for pups emerging from their natal burrow. Litter emergence dates were first precisely recorded (i.e. within 24 hours) in 1994. Pups were measured at emergence for mass (g) and anogenital distance (mm). Maternity was tentatively assigned based on the location of the emerging litter within the home range of an adult female late in lactation; maternity was then often confirmed by trapping the putative mother with her pups, and observations of affiliative interactions between the putative mother and pups (Ferron 1985). Most maternities were also confirmed by molecular parentage assignment (see below, Wells et al. 2017). Litters for which maternity could not be confirmed were excluded from this dataset. Adult females in this population live for an average of 2 years (Hostetler et al. 2012) and typically produce one or two litters; the oldest GMGS in this study was 9 years old and produced 7 litters.

### Variables

We define reproductive phenology as the timing of a female’s litter emerging from their natal burrow, at weaning. Female ground squirrels typically enter estrous and conceive a litter within a few days of emergence from hibernation (Williams et al. 2014, Morton & Gallup 1975), but may delay mating if they need to regain body condition before allocating energy to offspring (Millesi et al. 1999). We defined *absolute date of weaning* as the calendar date on which at least one pup was observed at or above a burrow opening in the mother’s home range. We defined *relative date of weaning* as the number of days after the date of complete snowmelt (0cm on ground) that pups emerged (i.e. absolute date of weaning minus date of snowmelt). Averages are given ± 1 S.E.

### Environmental tracking and potential limit to population plasticity

The Southern Rockies have experienced warmer and drier conditions over the past several decades, resulting in earlier spring snowmelt (Wadgymar et al. 2018, Cordes et al. 2020). These conditions have advanced flowering phenology of high-alpine species (Wadgymar et al. 2018) and likely the spring phenology of hibernating vertebrates in the same environments (Inouye et al. 2000). A directional relationship between phenology and time over the same period as environmental warming is consistent with climate-induced shift, but alone is not sufficient evidence as other non-climatic environmental factors may change over the same time period (e.g. resource availability, Lane et al. 2018). Hence studies need to examine the direct relationship between a climate-related cue (e.g. snowmelt) and phenology. Here we used a linear model to predict absolute date of weaning by date of complete snowmelt. Additionally, to test for a threshold response to earlier springs, we used the *segmented* package (Muggeo 2022) to evaluate linear vs nonlinear model fit (Iler et al. 2013). We used AIC ≥ 2 to distinguish if models had different fits to the data, with the lower AIC value indicating the better fit. All analyses were conducted in R v4.3.1.

### Repeatability of reproductive phenology

We calculated individual repeatability of litter emergence using the method of Lessels & Boag (1987). We excluded any litters for which emergence timing was uncertain (i.e. >48 hours since mother’s home range was last searched for emerging pups). We calculated repeatability first for *absolute date of weaning*, and then for *relative date of weaning*.

We expected that absolute date of weaning would not show significant repeatability, as squirrels are likely to adjust phenology to environmental conditions that cue resource availability such as snowmelt (Van Vuren & Armitage 1991; Prather et al. 2023), and inter-annual variation in date of snowmelt is high (Cordes et al. 2020). Instead, we expected that relative date of weaning would show significant repeatability, due to between-individual variation in sensitivity to environmental cues (e.g. Oosthuizen et al. 2023): for example, some females – perhaps those who regularly emerge from hibernation in good condition – may tend to produce litters soon after snowmelt, and other females may tend to produce litters substantially after snowmelt.

We also tested whether female age influenced reproductive phenology: we expected that older adults (2+ years) would wean litters earlier in the season than yearling (1 year old) females as older adults have finished somatic growth and typically emerge from hibernation in better body condition than yearlings (Dark et al. 1989; Howland et al. 2024). Age was known exactly for females who were born within the study area, and for those who immigrated into the study area in their natal summer and could be identified as juveniles based on their body mass at first capture (Howland et al. 2024).

### Heritability of reproductive phenology Matriline-only pedigree

Multilocus microsatellite genotypes were available for many individuals from 1996-2015 (Wells et al. 2017), and were used to confirm maternity for pups born 1996-2005 and 2007-2015. Candidate fathers (i.e. adult males) were not completely sampled in 1996-2005, precluding the consistent identification of unique fathers necessary for pedigree assignment. More complete parentage information with assigned paternities was available for 2008-2015 (Wells et al. 2017). However, as this represents a small subset of the dataset, we chose to construct a matriline-only pedigree (e.g. Reale 1999), in which we have high confidence of assignment.

### Animal model

We used an animal model to partition the variance in timing of litter weaning into genetic and environmental components (Wilson et al. 2010), using the package *MCMCglmm* (Hadfield 2010). Our animal model predicted log-transformed *weaning date* with no fixed effects (∼1) and female ID fit as a random effect to relate individuals to their additive genetic values through the matrilineal pedigree described above (Wilson et al. 2010). We used parameter expanded priors to improve chain behavior for variance components near zero: (Hadfield 2019): V=1, nu=0.002, alpha.mu=0, alpha.V=1000. We ran the model for 700,000 iterations, with a burn-in of 70,000 iterations and thinning interval of 700 to reduce autocorrelation. Heritability (h^2^) was calculated as V_a_/V_a_+V_r._ Variance components and heritability are presented as posterior modes with 95% highest posterior density intervals (Wilson et al. 2010).

Classically, heritability has also been estimated as the slope of the line of a parent-offspring regression (Falconer & MacKay 1996; Plard et al. 2014). To validate our animal model results with this method, we also fit a linear model, regressing a female’s first date of litter weaning (typically at age 1 or age 2) on her mother’s first date of litter weaning (typically at age 1 or age 2). To include weaning dates from multiple daughters of the same mother, we fit a multi-level model of daughter weaning date with mother weaning date as a fixed effect and mother identity specified as a random effect.

### Match-mismatch with vegetation phenology

We defined the onset of vegetation phenology as the first date of bare ground in spring (Van Vuren & Armitage 1991). We calculated the number of days between date of bare ground and a female’s litter emergence each year as the relative emergence date, and predicted this by the date of bare ground. This approach will necessarily produce a high level of autocorrelation, but here we are not seeking to explain the amount of variance (as it will be artificially high) but to examine the slope of this relationship. A flat line would indicate perfect plasticity in litter weaning phenology, in that females maintain a specific number of days between bare ground and litter weaning; a negative slope would indicate that as snowmelt advanced, the period between bare ground and litter weaning lengthened.

## Results

### Summary

We identified *N*=192 litters with certain weaning date from 1994 to 2023; these represented 1 to 6 litters from 104 unique mothers. Date of snowmelt ranged from April 23 at the earliest (in 2012) to June 19 at the latest (1995), with an average date of May 17 ± 12.8 days. Absolute dates of litter weaning ranged from June 8-August 11 (x=July 9 ± 0.79 days), and relative dates of litter weaning ranged from 19-81 days after snowmelt (x=51.8 ± 0.78 days). Average litter weaning dates advanced over the study period at the rate of 2.3 ± 0.9 days per decade (β_year_=-0.23 ± 0.09, t= -2.64, p<0.01; R^2^=0.04).

The relationship between date of snowmelt and litter weaning dates each year was positive (β=0.51 ± 0.05, t= 10.44, p<0.001; R^2^=0.36; Fig 1.): litters were weaned earlier in years of earlier snowmelt. A linear model for this relationship was supported over a non-linear model (AIC_linear_=834.92 vs. AIC_nonlinear_=838.11, ΔAIC= -3.19), suggesting no flattening of the response in the earliest springs.

**Figure 1.**
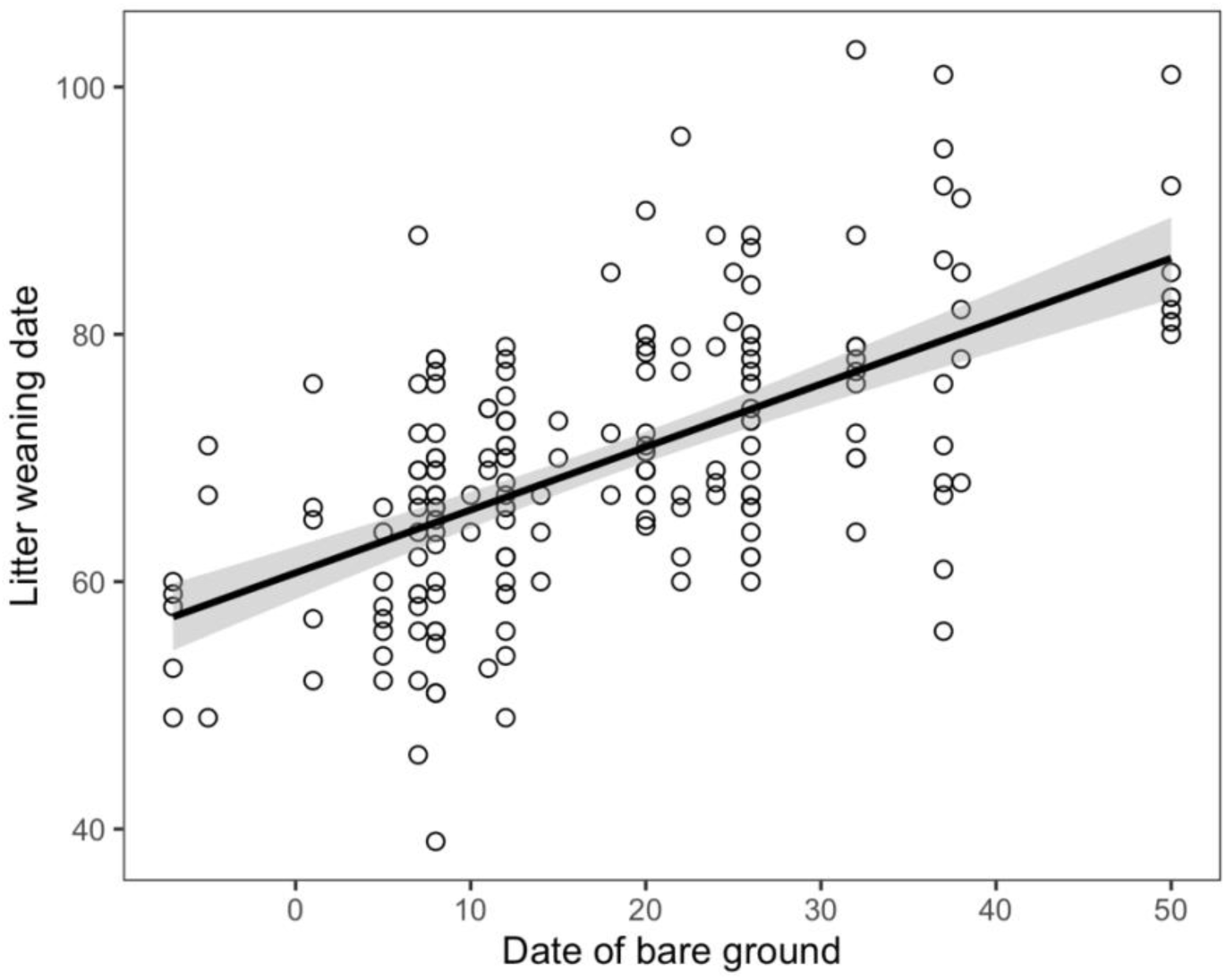
Individual dates of golden-mantled ground squirrel litter emergence each year from 1994-2023, regressed against date of bare ground that year. Dates are in number of days since April 30.

### Repeatability of reproductive phenology

Restricting the dataset to females with 2 or more litter weaning dates, we identified 141 litters belonging to 53 females (x = 2.66 litters/female: 32 females with 2 litters, 13 with 3 litters, 4 with 4 litters, 2 with 5 litters, and 2 with 6 litters). Absolute weaning date was significantly repeatable within female (r = 0.18, F_52,88_=1.58, p=0.03). By contrast, relative weaning date was not significantly repeatable (r = -0.12, F_52,88_ = 0.73, p=0.90). Female age affected the timing of litter weaning, as yearling females weaned litters significantly later than adult females 2 years or older (July 14 ± 1.6 days, vs July 6 ± 0.9 days; β= -6.63 ± 1.7, t= -3.82 , p<0.001; Fig2a). Additionally, individual females tended to shift litter weaning to earlier in the year between yearling and older ages (x = 5.82 ± 2.45 days earlier, N=24, Fig 2b).

**Figure 2.**
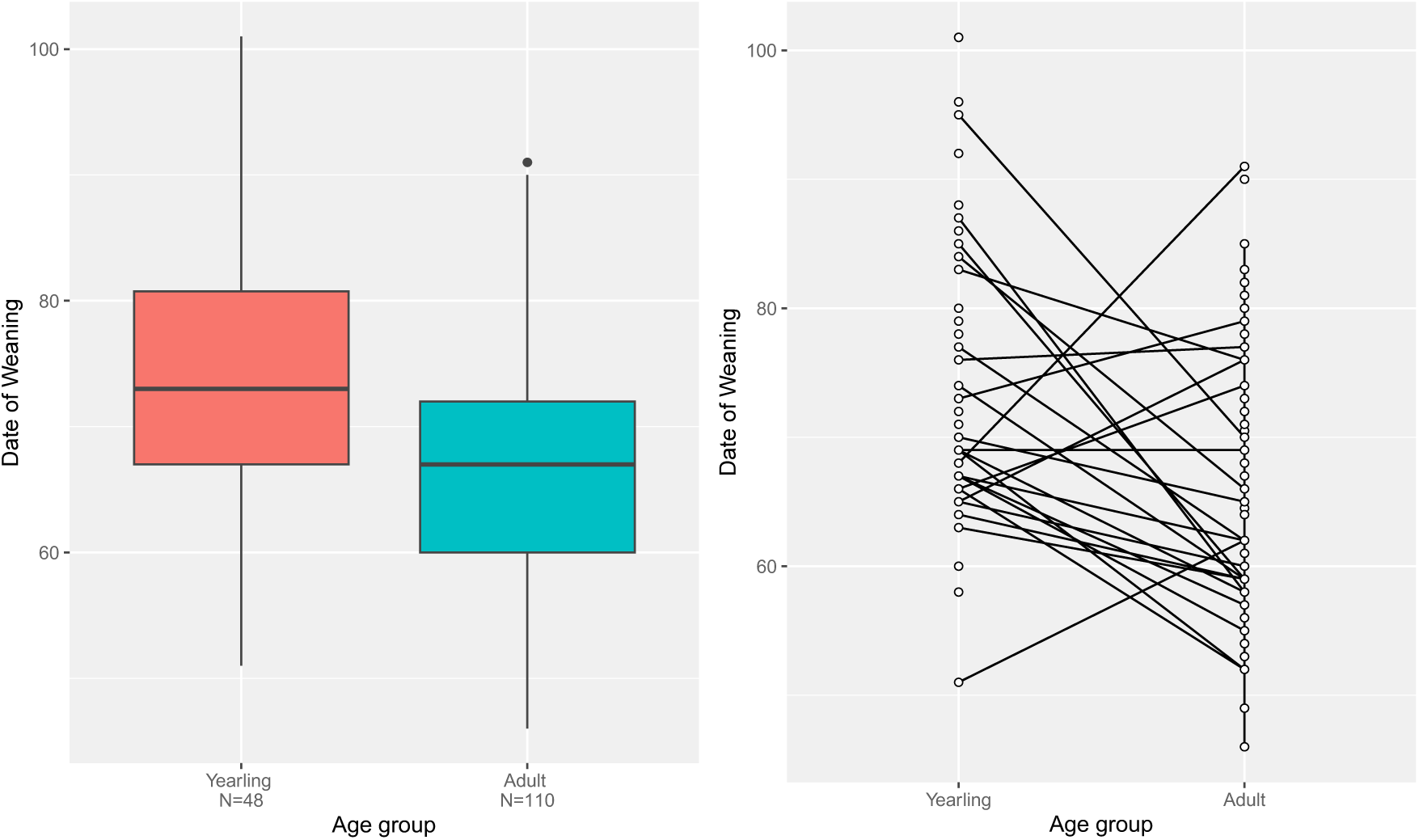
Differences in weaning dates by age group (a) at the population level, and (b) at the individual level. Open circles in (b) represent individual litter weaning dates; lines represent individual reaction norms between litters produced as a yearling (age=1) versus as an adult (age=2+).

### Heritability of reproductive phenology

Our maternal pedigree included 643 females, 123 dams, and 530 maternal links. The maximum number of generations in one matriline was 7. Of 104 unique females with litter weaning dates, 75 had known maternity and hence could be related to relatives via the pedigree. Autocorrelation was <0.06 for all lagged iterations beyond 0 (Lag 700, 3500, 7000, and 35000). The variance component for animal was near zero (V_a_=0.0008 [95% HDPI 0.0002-0.0108]) and smaller than the residual variance (V_r_=0.0220 [95% HDPI 0.0123-0.0320]). The posterior mode for heritability in timing of litter weaning was h^2^ =0.0362 (HDPI 0.0076-0.3763). When we ran the model again with a strong prior (V=1, nu=1), estimated heritability was extremely similar: mode h^2^= 0.0396 (95% HDPI 0.0086-0.3835).

We identified 69 mother-daughter pairs with first litter weaning dates for the parent-offspring regression. Maternal litter weaning date did not significantly predict daughter weaning date in either the linear (β=-0.06 ± 0.12, t= -0.52, p=0.61; R^2^=-0.01; Fig 3) or the multi-level model (β=-0.14 ± 0.14, t= -1.01), supporting the animal model mode of near-zero heritability of litter weaning date.

**Figure 3.**
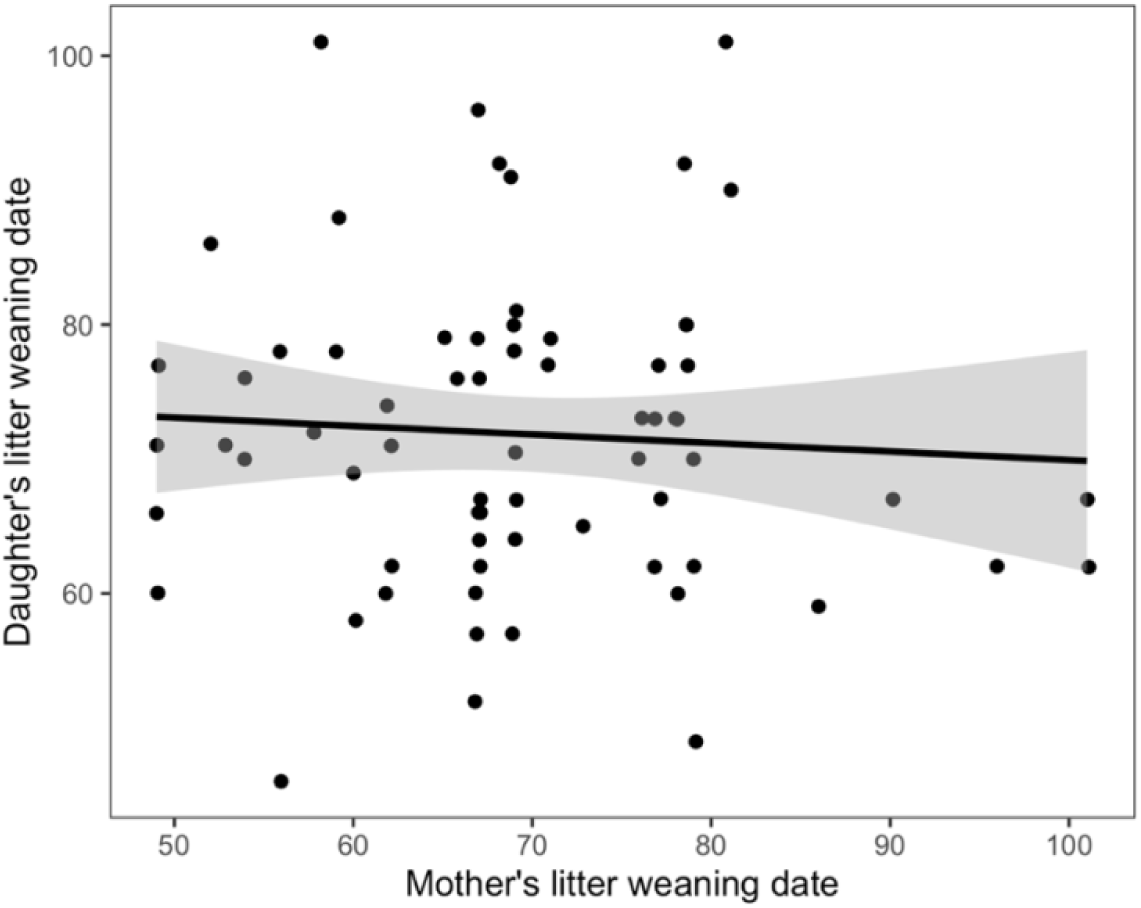
Regression of daughter’s first litter weaning date (in number of days since April 30) on her mother’s first litter weaning date, in a single population of golden-mantled ground squirrels (*Callospermophilus lateralis*) followed from 1995-2023 in Colorado, USA. Zero slope suggests no heritability in this measure of reproductive phenology.

### Match-mismatch with vegetation phenology

The relationship between relative litter emergence date and date of snowmelt was negative, with a moderate slope (β=-0.49 ± 0.05, t= -10.09, p<0.001; R^2^=0.35; Fig. 4). In the year of latest snowmelt, litters emerged an average of 35 days after the onset of vegetative growth, versus an average of 64 days in the years of earliest snowmelt (Fig 4).

**Figure 4.**
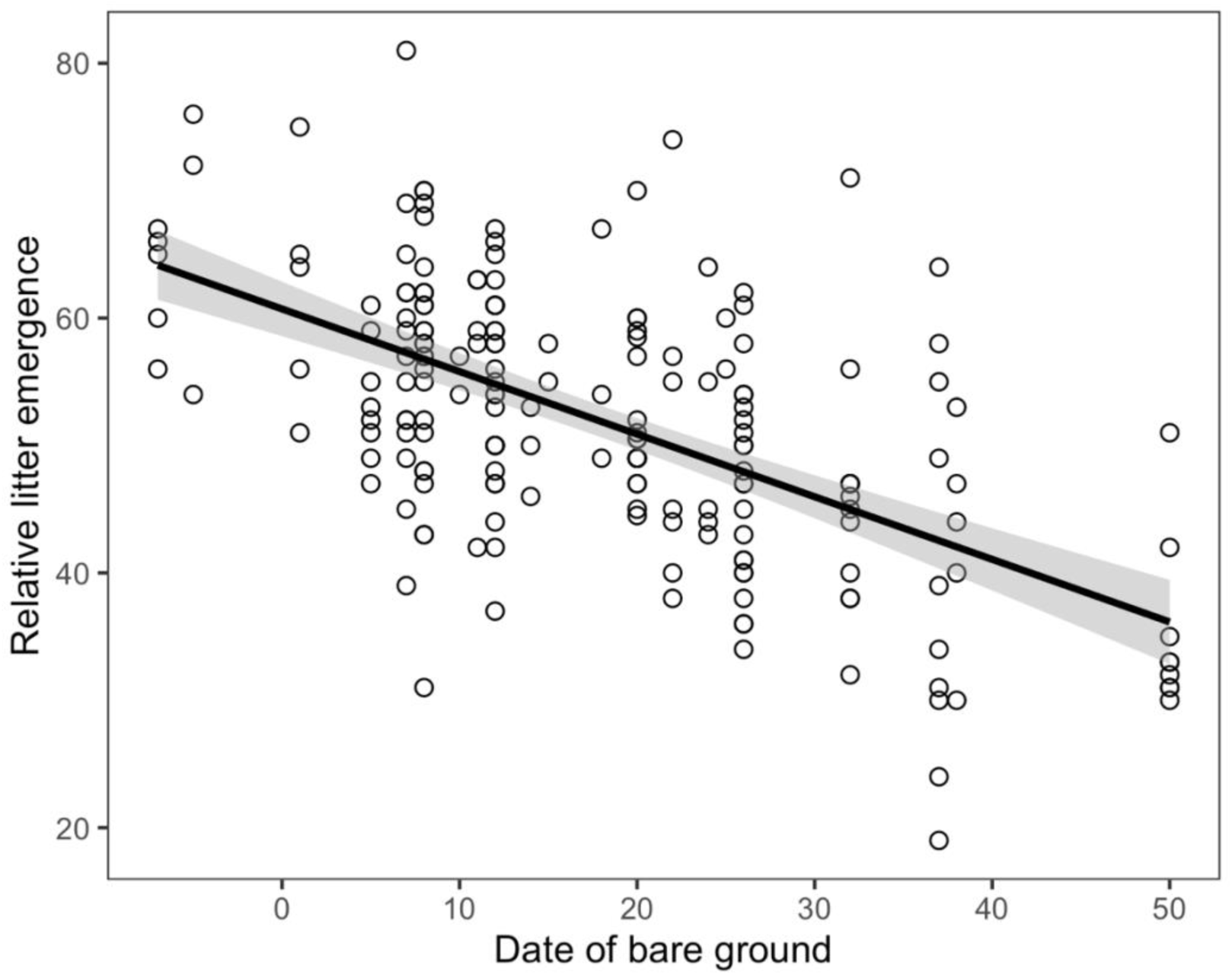
Weaning dates of individual golden-mantled ground squirrel litters relative to snowmelt (Date of bare ground=0), calculated as the date of litter emergence aboveground minus the date of bare ground that spring. Open circles indicate individual litters. Date of bare ground is reported as the number of days after April 30.

## Discussion

In this high-elevation hibernating mammal, we identified high plasticity in reproductive timing, in which litter emergence at the population level has been closely tracking variable spring snowmelt. Over time, average litter emergence has advanced 2.3 days/decade with earlier springs, with no sign of a threshold in response at the earliest of spring conditions. However, litters born in early springs emerge relatively later in the vegetation phenological cycle than those born in late springs, raising the possibility of trophic mis-match. At the individual level, reproductive phenology was repeatable but not heritable, suggesting little scope for microevolution in this trait. Population-level plasticity despite individual repeatability may point to the importance of external or internal cues during development that set the initial timing of circannual cycles (e.g. Hazlerigg & Lincoln 2011), or the contribution of new recruits to population change (e.g. Gill et al. 2014).

Snowmelt date has advanced significantly at our study site over the preceding decades (Cordes et al. 2020), and plasticity in reproductive phenology of the golden-mantled ground squirrel appears to be tracking this change. The rate of advancement in reproductive phenology is comparable to that observed for arctic seabirds (1.9days/decade, Sauve et al. 2019) and another hibernating ground squirrel (1.7days/decade, Ozgul et al. 2010), but slower than that observed for several ungulates (4.2 days/decade, Moyes et al. 2011; 5.8days/decade, Renaud et al. 2019,); like ice-obligate seabirds in the high Arctic, high-elevation hibernating squirrels are likely constrained by short breeding seasons that begin with snowmelt, versus ungulates with longer gestation periods that begin the previous fall. Importantly, a linear relationship between litter weaning and spring snowmelt was supported over a nonlinear (i.e. threshold) model, suggesting that the timing of litter weaning may continue to advance should spring snowmelt conditions begin even earlier with warming and drying conditions. Despite the insight threshold models have yielded for flowering phenology (e.g. Iler et al. 2013), we were not able to find other examples where they had been tested for vertebrates.

Plant species at high elevations initiate growth and reproduction in a predictable sequence each year (CaraDonna et al. 2014) beginning with bare ground in spring and ending with hard frost in fall (Van Vuren & Armitage 1991). Hence, the species composition of plants available to herbivores varies with the number of days since snowmelt. For a generalist species like the golden-mantled ground squirrel, many plant species are consumable (Olin 1961, Carleton 1966). However, the amount of vegetation and specific nutrients available (e.g. in growing vegetative vs. reproductive structures) depends on timing in the sequence of vegetative phenology. Additionally, the nutritional quality of vegetation generally declines over the summer, as lignin content increases (Klein 1990). While litters emerge later in relation to the vegetative phenology in years with early springs, they still emerge earlier in the calendar year. More days to put on mass for hibernation may offset the reduction in vegetation quality (Howland et al. 2024). However, years of early snowmelt are also associated with lower precipitation, leading to foresummer drought and lower primary productivity (Sloat et al. 2015), making early springs a potential double-negative for growing pups or females seeking to regain mass before hibernation. In future, individual growth data are needed to evaluate this potential mismatch, and to add to the scant analyses on trophic mismatch in terrestrial mammals (Kharouba & Wolkovich 2023).

Individual dates of litter weaning were significantly repeatable to calendar date (absolute timing), but not number of days since snowmelt (relative timing). The repeatability to calendar date suggests primacy of endogenous circannual cycles over one external cue – the onset of vegetative growth – and is consistent with observations of circannual cycles in captive golden-mantled ground squirrels that were insensitive to changes in photoperiod and food availability (Pengelley & Fisher 1957, 1963; Pengelley & Asmundson 1974). However, external cues other than vegetative growth – such as temperature in fall (Prather et al. 2023) or during hibernation (Joy & Mrosovsky 1983) – may be essential to entrain the circannual cycle of this species, which once set continues through the reproductive lifespan of an individual squirrel. Future work should seek to evaluate and identify these cues.

Despite individual repeatability, reproductive phenology showed near-zero heritability in this small hibernator. Longer-lived organisms are likely to experience more variation over their lifetimes, and hence are expected to show more plasticity and less heritability than shorter-lived organisms (Chmura et al. 2019). Yet estimated heritability in hibernation emergence is higher for the shorter-lived Colombian ground squirrel (*h^2^*=0.22 ± 0.05, Lane et al. 2011) vs. the longer-lived yellow-bellied marmot (*h^2^*=0.16, 95% HDPI 0.10/10.26, Edic et al. 2020); both are higher than the estimated heritability of reproductive phenology in the golden-mantled ground squirrel (this study) or yellow-bellied marmots (*h^2^=* 0.05 ± 0.06, St. Lawrence et al. 2024). Oestrous dates are highly genetically correlated with emergence dates in some hibernators (Lane et al. 2011), but vary substantially in others (St. Lawrence et al. 2024). The annual cycles of most animals include multiple phenological events; if reproductive phenology is less heritable than and relies on different cues than migration or hibernation, mistiming between those phases may become increasingly common (Visser & Gienapp 2019).

What are the likely consequences of low heritability in reproductive phenology? Precision in the estimate of heritability in the animal model was limited by our sample size: while the posterior mode was very small (0.03), the HDPI was wide. Yet the daughter-mother regression indicated little heritability of reproductive timing, and therefore little potential for microevolution of this trait (Wilson et al. 2010). Micro-evolutionary responses are difficult to detect in wild populations, but at least partially underlie phenological shifts in spring migration in the common tern (*Sterna hirundo*, Moiron et al. 2024), and advancing parturition date in red deer (*Cervus elaphus*, Bonnet et al. 2019). For ground squirrels, the low magnitude of heritability suggests that selection will not be able to optimize adaptation to environmental change and facilitate long-term persistence in the face of directional change (Charmantier & Gienapp 2014). However, substantial plasticity in reproductive timing suggests that the changing environment will not select against specific genotypes and systematically reduce variation in populations exposed to earlier springs; given that climate variability is another challenge to vertebrate adaptation (Clark-Wolf et al. 2024), plasticity in reproductive timing is likely adaptive in an environment with high inter-annual variability in snowmelt. As we detected no threshold in plasticity in the earliest spring conditions, phenotypic plasticity at the population level appears currently sufficient for short-term adaptation to varying conditions.

But how do we reconcile our findings that individual reproductive phenology was significantly repeatable to calendar date, yet at the population level reproductive phenology was tracking snowmelt, which was highly variable among years? We consider at least three possible explanations. First, despite significant repeatability to calendar date, the magnitude of repeatability was low: reproductive phenology was still mostly plastic within individuals (>80%). Second, despite individual repeatability of experienced breeders, new recruits may be tracking environmental conditions and driving the population-level plasticity as has been found for migratory godwits (Gill et al. 2014). Given the often short lives of adult reproductive females in golden-mantled ground squirrels (average=2 years, Hostetler et al. 2012), approximately half of the females in our dataset reproduced just once (51/104=49%) and that reproductive bout may have been more likely to reflect that year’s snowmelt conditions. For repeat breeders, timing of the first reproductive bout may set a female’s individual circannual cycle to cyclical patterns of tissue histogenesis (Hazlerigg & Lincoln 2011), modified by her body condition at the end of hibernation; this would explain the small advances in reproductive phenology of adults versus yearlings (this study), as ground squirrel adults 2 years and older are typically in better body condition than yearlings in spring (Howland et al. 2024). Last, selection against reproductive females who emerged too early – either via mortality or litter loss during gestation or lactation – may shift the average reproductive phenology of *successful* litters to match snowmelt conditions. If this were the case, then population-level plasticity that appears adaptive may come at the expense of phenologically-mismatched individuals, potentially reducing population size and long-term persistence. Tracking the selective disappearance or reproductive failure of early-emerging females would be a valuable future direction to evaluate this potential mechanism.

Assessment of phenological shifts in mammals has lagged behind that of other taxa, particularly birds (Radchuk et al. 2019). Where studied, the ability to track changes in spring phenology has been associated with higher fitness (Cleland et al. 2012), but the rate of phenological adaptation may not be fast enough for long-term persistence (Radchuk et al. 2019). Moreover, shifting phenology has important consequences for conservation planning, but needs to be considered more consistently (Ettinger et al. 2022). In addition to linking phenological shifts to fitness outcomes, identifying which species have reached a threshold of plasticity to changing conditions and which have not will be valuable for directing conservation attention.

## Supplemental Figures

**Figure S1.**
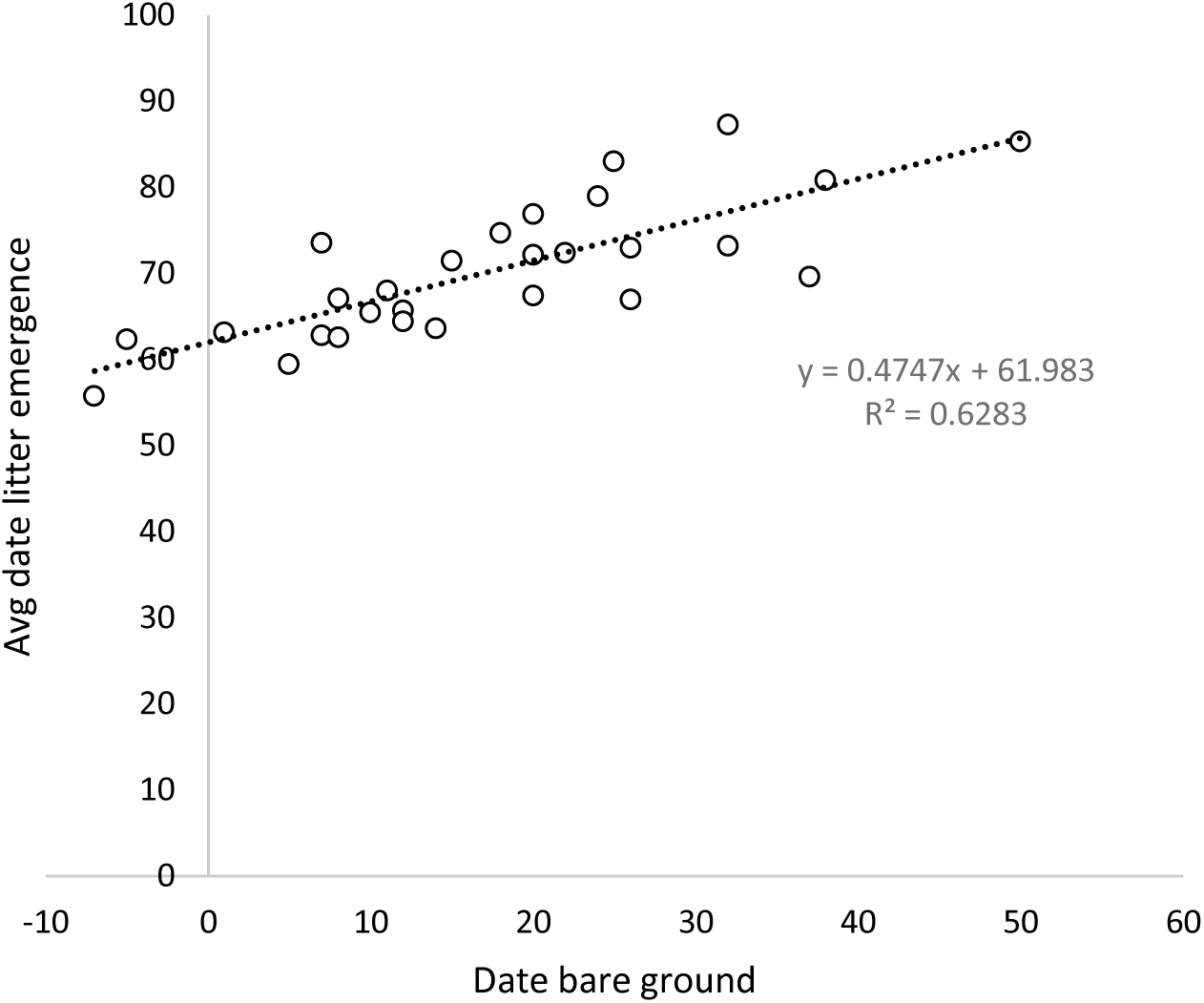
Average date of golden-mantled ground squirrel litter emergence each year from 1994-2021, regressed against date of bare ground that year. Dates are in days since April 30.

**Fig S2.**
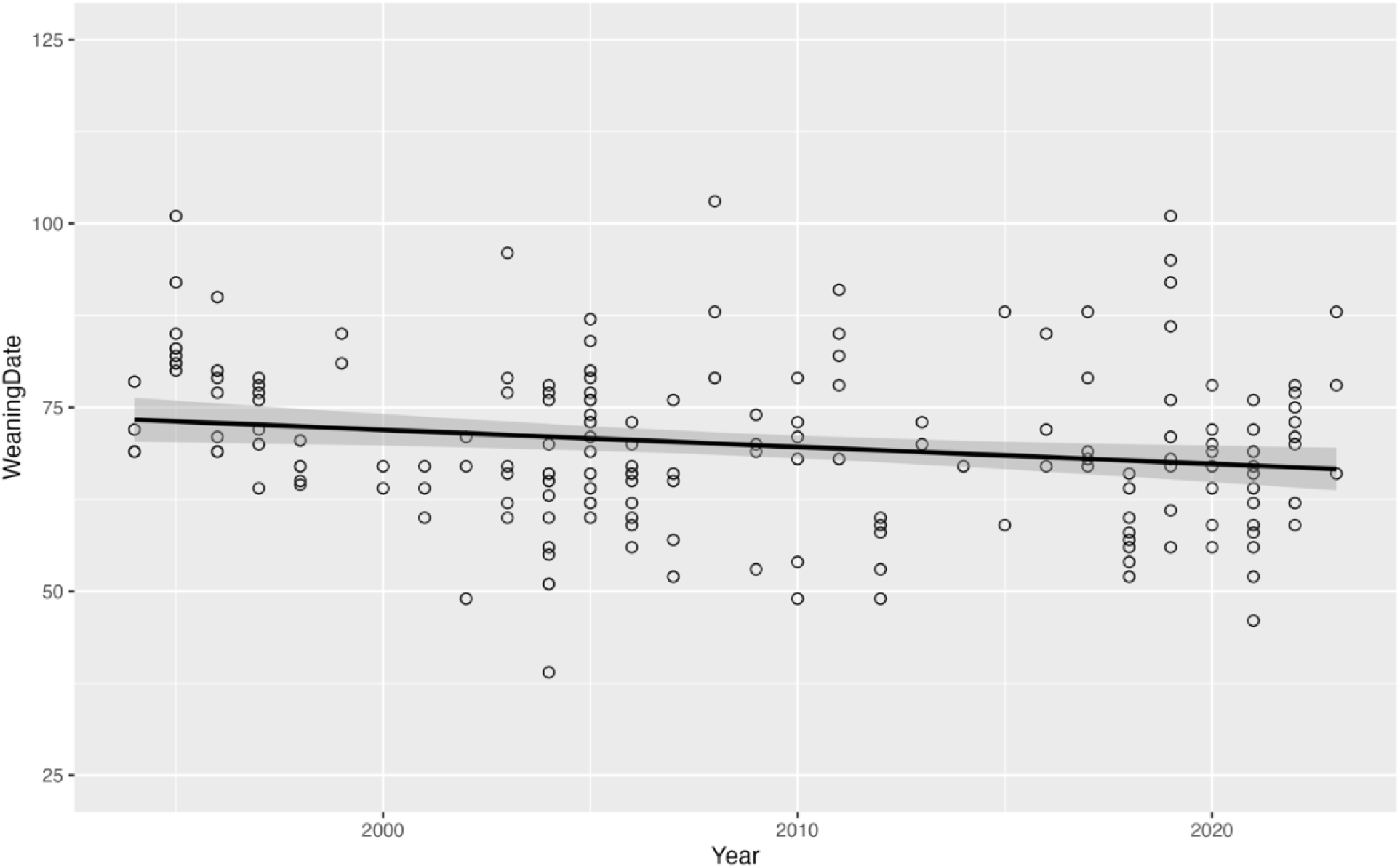
Litter weaning dates over time, from 1994-2023. Dates are in days since April 30. Fitted line indicates significant advancement over time of -0.23 days/year (B_year_=-0.23±0.09, t=-2.64, p=0.009).

